# Systematically understanding the key roles of microglia in AD development

**DOI:** 10.1101/601054

**Authors:** Zhiwei Ji, Changan Liu, Weiling Zhao, Claudio Soto, Xiaobo Zhou

**Author notes:** Correspondence to: Xiaobo Zhou,.

## Abstract

Alzheimer’s disease (AD) is the leading cause of age-related dementia, affecting over 5 million people in the United States. Unfortunately, current therapies are largely palliative and several potential drug candidates have failed in late-stage clinical trials. Studies suggest that microglia-mediated neuroinflammation might be responsible for the failures of various therapies. Microglia contribute to Aβ clearance in the early stage of neurodegeneration and may contribute to AD development at the late stage by releasing pro-inflammatory cytokines. However, the activation profile and phenotypic changes of microglia during the development of AD are poorly understood. To systematically understand the key role of microglia in AD progression and predict the optimal therapeutic strategy *in silico*, we developed a 3D multi-scale model of AD (MSMAD) by integrating multi-level experimental data, to manipulate the neurodegeneration in a simulated system. Based on our analysis, we revealed how TREM2-related signal transduction leads to an imbalance in the activation of different microglia phenotypes, thereby promoting AD development. Our MSMAD model also provides an optimal treatment strategy for improving the outcome of AD treatment.

## Introduction

Alzheimer’s disease (AD) is one of the most significant public health problems of the 21st century. It is estimated that the number of AD patients in the USA will increase from 5 million to 15 million by 2050, and the annual cost of care is projected to reach $1.1 trillion [1-3]. Unfortunately, current treatments of AD can only serve to alleviate symptoms in a short period, and there is no cure for this disease or a way to stop or slow its progression [4-6]. It is due to an incomplete understanding of the biological mechanisms underlying its pathogenesis on the molecular, cellular, and tissue levels [7]. Hence, there is an urgent need to improve our understanding of the molecular mechanisms that drive the development of late-onset AD.

The classical hallmarks of AD pathology are the accumulation of extracellular amyloid plaques and intracellular neurofibrillary tangles [8, 9]. Dysregulated amyloid beta metabolism has also been shown to promote insulin resistance in AD [10, 11]. One of the most striking hallmarks of AD is microglia-mediated neuroinflammation [12]. Microglia, the tissue-resident macrophages in the brain, are damage sensors of neurodegeneration in the AD progression [13]. Microglial activation can be categorized into two opposite types: pro-inflammatory M1 phenotype (neurodegenerative) and anti-inflammatory M2 phenotype (neuroprotective) [14, 15]. It is nowadays accepted that there is a dynamic microglia turnover in the brain and that microglia phenotype may change depending on aging or the stage of the disease [14]. However, the activation profile and phenotypic changes of microglia during the development of AD are poorly understood. Therefore, understanding the sequential and timing-associated changes in M1/M2 activation and the potential factors/pathways that control microglia activation may provide better therapeutic benefit.

Currently, genome-wide association studies (GWAS) in AD have uncovered the enriched genes in microglia, such as CD33, CR1, EPHA1 and TREM2 (triggering receptor expressed on myeloid cells 2) [16]. Recent studies found that TREM2 increased AD risk by about 3-times [17, 18]. TREM2 and its adaptor DAP12 exert neuroprotective effect in microglia by mediating microglial survival and Aβ clearance [19]. However, sTREM2, the soluble fragment of TREM2, is strongly correlated with amyloidosis and microglial activation, suggesting that sTREM2 serves as a biomarker for triggering inflammation response via M2/M1 switch [20]. The definitive mechanism of TREM2 and sTREM2 in AD remains largely unknown [21]. Integrating large-scale genomics data from public databases (e.g. ADNI [22], GEO, *etc*) will be helpful in understanding how TREM2 and sTREM2 differentially modulate microglia phenotype activation during AD progression.

In the present study, we integrated a set of large-scale genomics data and identified the genes and pathways associated with the phenotypic heterogeneity of microglia cells. Our analysis showed the transcriptional programs in microglial cells over time. At the early AD stage, the major changes in microglia were characterized by up-regulation of TREM2/DAP12, phagocytosis-related genes, and anti-inflammatory genes. Elevated expression of sTREM2/NFKB and pro-inflammatory genes were observed at the late stage. To systematically understand TREM2-regulated imbalanced activation of M1/M2 microglia leading to AD progression and predict the optimal therapeutic strategy, we further developed a predictive Multi-scale Model of AD (MSMAD) by integrating multi-level experimental data. After parameter tuning, the outcomes of our model under different contexts fit the experimental observations well. Finally, we used the MSMAD model to predict the effect of single or combined treatments. Our simulation indicates that switching microglia activation toward the M2 phenotypes and activating insulin metabolism appear to impair AD development. In summary, this study revealed the key cytokines/pathways-induced microglia activation during AD progression and also provide an optimal therapeutic strategy for improving the outcomes of AD treatment.

## Results

### Identifying temporal changes of intracellular pathways in microglia in AD

To explore the phenotypic heterogeneity of microglia in response to neurodegeneration, we selected three datasets from GEO to identify the transcriptional programs in microglia cells over times. Firstly, we analyzed the dataset GSE103334 generated from microglia isolated from hippocampus tissues in healthy (CK control) and AD mouse (CK-p25 transgenic [23]). At 2 weeks after p25 induction, CK-p25 mice exhibit DNA damage and increased amyloid-β level, followed by progressive neuronal and synaptic loss with cognitive impairment, which is more severe by 6 weeks. The genes in hippocampal microglia were profiled before p25 induction and 1, 2, and 6 weeks after p25 induction. Mathys, *et al*., demonstrated that the CK-p25 mouse model displays key pathological hallmarks of AD in a temporally predictable manner, to study the response of microglia at temporal- and single-cell resolution [24]. In our study, we screened out three clusters of cells to represent control (cluster 2), early response (cluster 3), and late response (cluster 6) of microglia, respectively. SCDE was used to determine the differential expression of genes between different groups [25]. **Fig 1A** presents the relative expression of Lgals3, CD68, B-catenin, and IGF1 in microglia, indicating the up-regulation of phagocytosis, anti-inflammatory effects, and cell survival in early stage of AD. **Fig 1B** shows that NFKB, TNF-α, IL-1 β in microglia are up-regulated in the late stage relative to normal control. CD68 level was reduced in late response group relative to early response one. Particularly, the expression of SYK in late stage was down-regulated. Moreover, we will validate the above molecular factors of CK-p25 mice with the experimental data from 5xFAD mice because previous study reported that the transcriptional profiles from these two types of mice show similar concordance with human AD brain signatures [23]. 5xFAD mouse model harbors five early-onset familial AD mutations and allows a comprehensive understanding of molecular events occurring in 6-9 months old 5xFAD mice during the early stage of AD [26-28]. Therefore, we further selected GSE65067 dataset to examine the role of Trem2 on the early response of microglia to Aβ deposition. Microglia were purified from 8.5-month-old WT and 5xFAD mice. Differentially expressed genes (DEGs) were screened with R limma package [29]. Here, we analyzed 83 representative genes (**Supplementary Data File 1**), including 67 DAM (disease-associated microglia [30]) genes [31], and 16 microglia-associated factors [31-33]. **Fig 1C** shows that Trem2, AKT, phagocytosis-related genes (Lgals3, and CD8), and anti-inflammatory factor IGF-1 are significantly up-regulated microglial cells. Finally, we used dataset GSE104775 to study the temporal changes of gene expression in microglia in the early stage of AD in 5xFAD mice (**Supplementary Data File 2**). As shown in [31], the gene expressions in 5xFAD have no significant changes at 2-month-old mice and Trem2, Tyrobp, SYK, CD68, Lgals3, and TGFB1 are markedly increased at 4- and 7-month-old 5xFAD mice (**Fig 1D**). The findings shown in **Fig 1C-D** are close to **Fig 1A-B**. Taken above together, our analysis indicated that TREM2/DAP2 signaling was activated in the early response, which may facilitate phagocytosis and induce microglia proliferation. As the downstream of Trem2 signaling, Wnt/β-catenin pathway further promote microglia cell survival (**Fig 1E**). However, the neuroprotective effects of microglia is lost in late-term AD due to the down-regulation of Trem2 signaling and activation of NFKB-mediated pro-inflammatory effects (**Fig 1F**). The accelerated neurodegeneration might be caused by the elevated sTREM2 concentration, which is released from Trem2 proteolytic cleavage [21].

**Fig 1.**
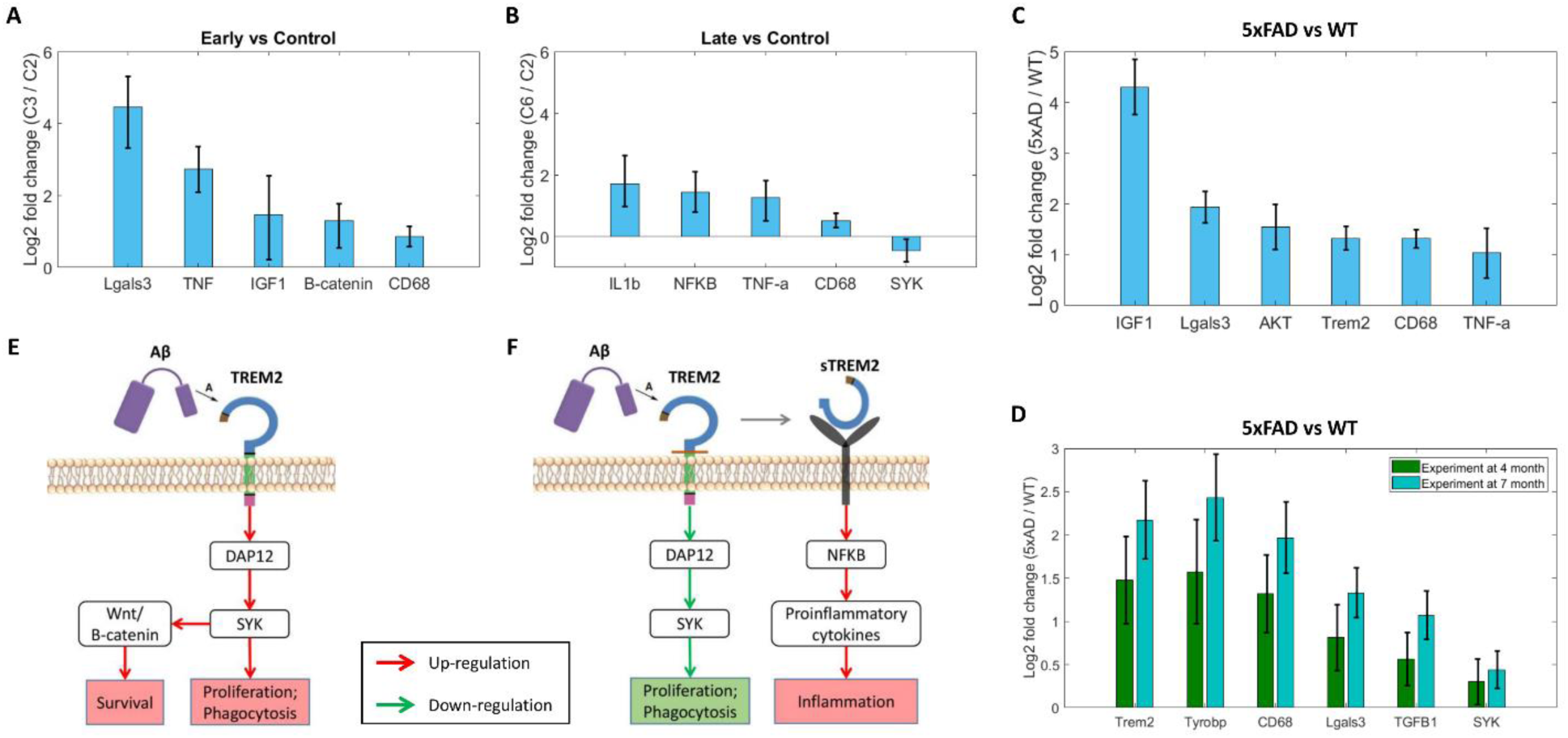
Identification the temporal changes of key factors in microglia cells. **A-B**. DEGs in the early and later response of microglia (GSE103334). C**-D**. DEGs identified by GSE65067 and GSE104775. **E-F**. Inferred intracellular pathways of microglia in early and late response. Red arrows: up-regulated pathways; Green arrows: down-regulated pathways.

### Inferring intracellular pathways of neurons in AD

To investigate the intracellular signaling pathways of neurons in the complicated brain microenvironment (mE), we selected a representative RNA-seq dataset (GSE75431) from GEO, which was generated from brain cortex in PS2APP AD mouse model. Three types of cells (astrocytes, microglia, and neurons) were isolated and sequenced. The gene expression profiles were collected from 7 and 13 months old PS2APP AD mice. By using limma package, we identified 802 and 1469 DEGs in the neurons from 7 and 13 months of mouse model, respectively (P-value<0.05). In the 7 months AD mouse model, the expressions of IGF1, Tyrobp, CSF1, and CD68 were up-regulated in microglial cells (FC > 1.5), and the expression of MAPK1 in neurons was significantly elevated. We also found the increase of GSK3β and decrease of PKC in neurons in 13 months AD mouse model relative to control (FC > 1.5). The details of differential expression analysis was shown in **Supplementary Data File 3**.

The inferred specific signaling pathways in neurons related with AD were shown in **Fig 2**, which include three pathways induced by three different ligands. IGF-1 signaling regulates neuronal growth, synaptic maintenance, neuroprotection via MAPK pathway and PI3K/AKT pathway [34]. Accumulation of Aβ oligomer may: 1) induce neurotoxicity by inhibiting the PI3K pathways in neuronal cells [35]; and 2) lead to increased TNFR, resulting in inhibitory phosphorylation of IRS-1/PI3K/AKT. The decreased brain insulin signaling leads to increased GSK-3β activity, causing increased abnormal tau phosphorylation [36]. The inferred pathways were used in the intracellular level of the 3D MSMAD model to stimulate the dynamics of signal transduction in response to extracellular microglia-neuron interactions.

**Fig 2.**
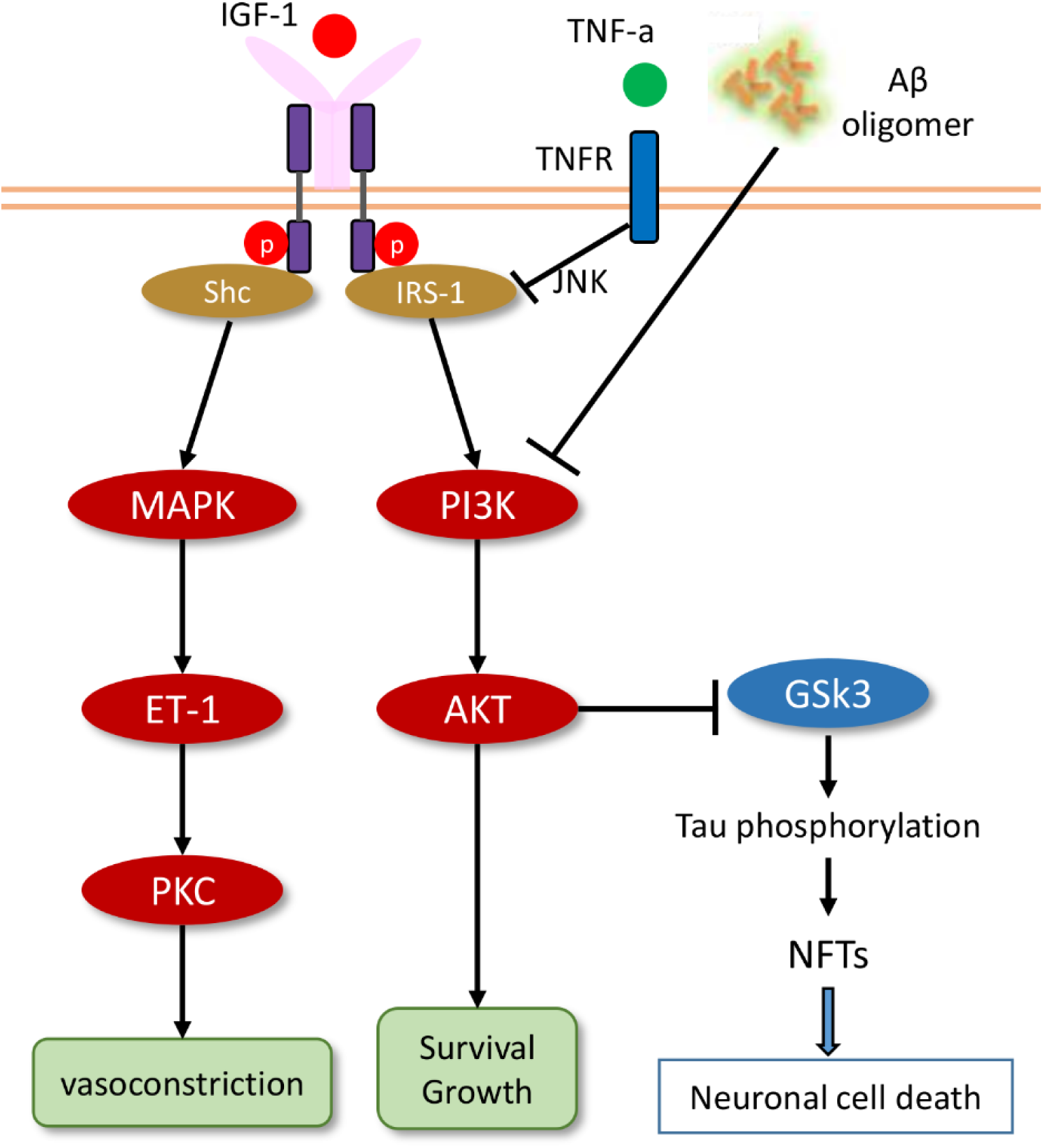
The inferred specific pathway of Neuron based on GEO dataset GSE75431.

### 3D Hybrid multi-scale modeling of the AD progression

#### Hypothesis

Based on the comprehensive analysis of the collected experimental data, we proposed the following scheme (**Fig 3**). Aβ peptides are secreted from neuron cells and accumulated to form Aβ oligomer and senile plaque in the early-onset of AD. The toxic effects of Aβ oligomer can induce neuron cell death. In another aspect, Aβ oligomer is able to attract and stimulate the resting microglial cell (M0 microglia) in the initial immune response. Consequently, the alternative activation (M2) of microglia cells leads to neuroprotection, because 1) the increased secretion of IGF-1 from M2 promotes neuronal cell survival and proliferation via PI3K/AKT and MAPK pathways and 2) activation of TREM2 signaling in M2 microglia induce Aβ oligomer clearance before formation of senile plaque. In the late-onset, M2 microglia gradually switch to M1 microglia via TREM2-related signaling, and then aggravate neurodegeneration. NFKB pathway was activated in M1 cells, which cause the generation of inflammatory factors (e.g. TNF-*α*, and IL-1β). Alterations of TREM2 signaling reduced the ability of M1 microglia to clear Aβ. Moreover, as the proportion of M1 microglia increases, the concentration of sTREM2 in CSF increases dramatically.

**Fig 3.**
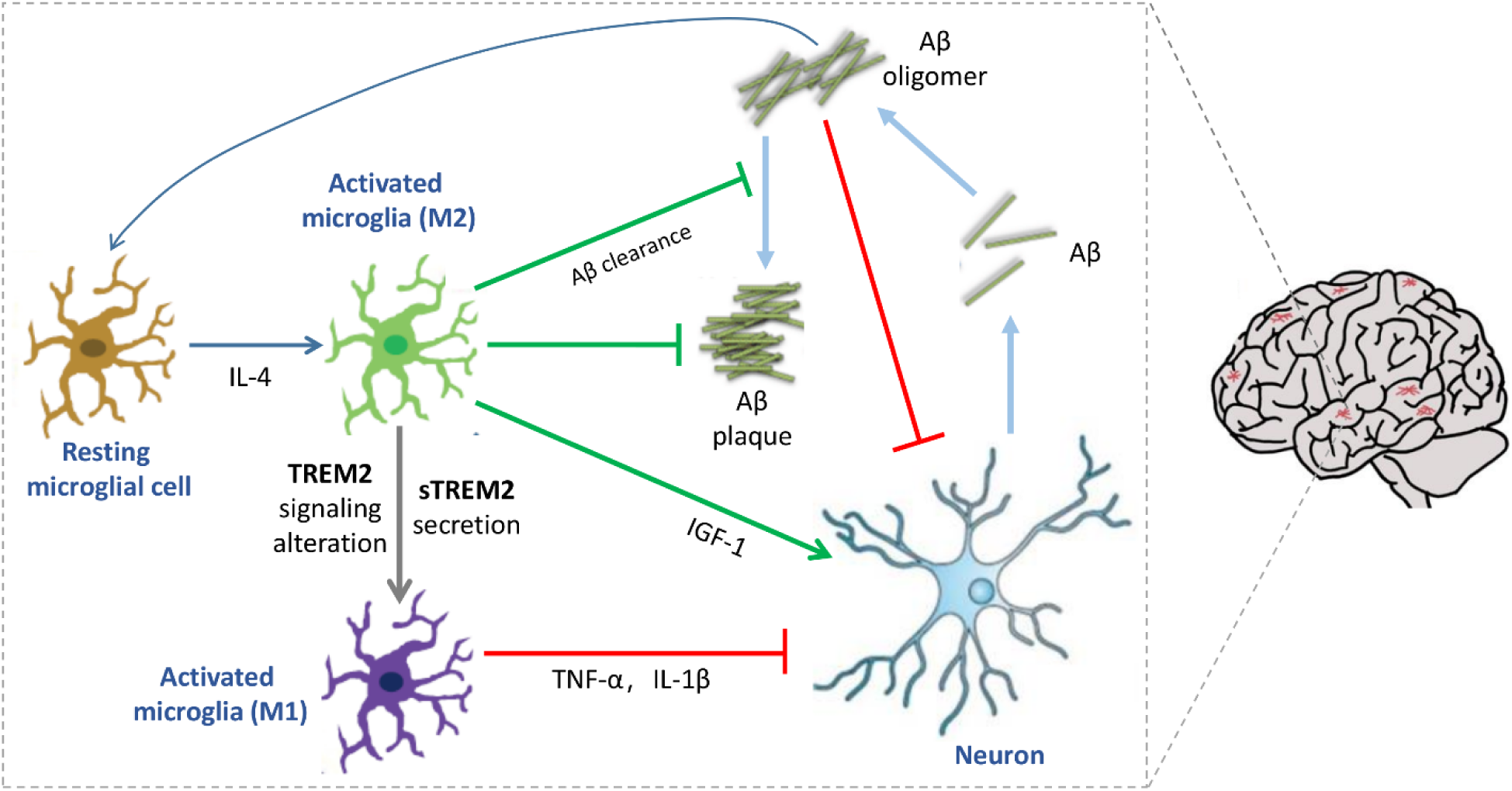
Systems diagram of MSMAD model.

#### Systems modeling

We developed a novel 3D multi-scale model of AD (MSMAD) to study the phenotypic heterogeneity of microglia activation during AD progression, as well as to verify the proposed the scheme showing in **Fig. 3**. The MSMAD model combines a 3D multi-scale agent-based model (ABM) for neurodegeneration and immune response, and a Hill function system for dynamic signaling transduction. Our MSMAD model defines three type of agents to represent neuronal and microglia cells, and Aβ. In addition, eight types of small molecules diffuse and degrade in the simulated mE, including Aβ from Neurons, IL-4 from M0 microglia, IGF-1 from M2 microglia, TNF-*α*, and IL-1β from M1 microglia, as well as Aβ oligomer and Aβ plaque accumulated from Aβ. AD progression is simulated at intracellular, intercellular, and tissue levels in our MSMAD model. Intracellular signal transduction was modeled by Hill functions. The input parameters of Hill functions are the local concentrations of the cytokines shown in **Fig 1-2**. The intercellular communication reflects the cell-cell interactions, including Ligand-receptor mediated cell-cell interactions, as well as physical interactions with the other cells. The tissue scale of our model reflects the AD brain tissue (e.g., hippocampus) and Aβ plaque formation via various intercellular cell-cell interactions. Moreover, the MSMAD model integrates a series biological events spatially and temporally. Spatially, the intracellular pathways are encapsulated into cells to determine the immune response or neuronal cell behavior (Aβ secretion, migration (**Fig S1**), proliferation or apoptosis). In turn, cytokines diffusing in 3D space trigger the intracellular signaling pathways via their receptors, resulting in the alteration of cell behaviors. Temporally, it covers signaling pathway dynamics (minutes to hours); cell division, apoptosis and local migration (hours to days); treatment responses and neuron growth (days to weeks). The cell behaviors and cytokine diffusion in 3D ABM model are updated every 2 hours. The details of the rules for agent-agent and agent-mE interactions are described in **Supplementary Information**.

#### Hybrid model implementation

We implemented the 3D multi-scale system (MSMAD) by incorporation of ABM and Hill Functions. The ABM is a stochastic model and Hill functions are combined as a simplified continuous model. The communications between them were mediated by cytokines/growth factors described above. We simulated the whole process for up to 6 weeks according to the time line in CK-p25 mice [24], and exported the dynamic profiles of all the cell populations and cytokines. Based on all the observed data collected from the literature (**Table 1**), we manually tunned all the parameters associated with ABM model in MSMAD for minimizing the errors of data fitting. All the parameters of ABM model are presented in **Table S1**. The framework of ABM was designed and achieved with C++. The whole model was implemented under Linux on the platform of TACC.

**Table 1.** The experimental data used for data fitting of MSMAD model.

|  | NC | AD | Models | PMID |
| --- | --- | --- | --- | --- |
| Neuron cells ($\times 10^6$) | $6.9 \pm 1.1$ | $3.6 \pm 1.6$ | AD patients | 8699259 |
| Microglia activation (FC) | 1 | 1.36 | AD patients | 28122877 |
| Plaque radius ($\mu\text{m}$) | $5.413 \pm 0.04$ | $13.41 \pm 1.65$ | APPPS1 mice | 22993126 |
| Microglia per plaque (FC) | 1 | 4.28 | 5xFAD mice | 25728668 |
| sTREM2 (pg/ml) | $195.5 \pm 45.1$ | $231.2 \pm 74.2$ | AD patients | 26754172 |
| TNF-α (pg/ml) | $20.56 \pm 10.90$ | $52.23 \pm 22.36$ | AD patients | 15936505 |
| IL-1β (FC) | 1 | 6.0 | AD patients | 2529544 |
| IGF1 (FC) | 1 | 0.3 | AD patients | 24301648 |

### Model evaluation and validation

To test the performance of our established model, *in silico* simulations under several contexts were evaluated using the experimental data collected from the literature (**Table 1**). Firstly, we simulated the whole process of AD development in mouse model from the initial state to 6 weeks. The dynamic changes of neuron and microglia population in the simulated mE were predicted. **Fig 4A** shows that neuron cell survival is reduced 0.47 folds and microglia activation (M1, and M2) is increased 0.34 in AD patients. **Fig 4B** represented that AD progression results in plaque accumulation and increased the numbers of microglia surrounding the amyloid plaque. The predicted results shown in **Fig 4A-B** are consistent with the experimental data reported in [15, 37-39]. Furthermore, we calculated the relative changes of cytokines and growth factors in AD relative to normal control. As shown in **Fig 4C-D**, the concentrations of sTREM2, TNF-α, and IL-1β are sharply increased, and IGF1 level is reduced. The simulation results are close to the previous findings reported in [40-43]. Taken together, our results indicate that the MSMAD model fits the observed data very well under different contexts.

**Fig 4.**
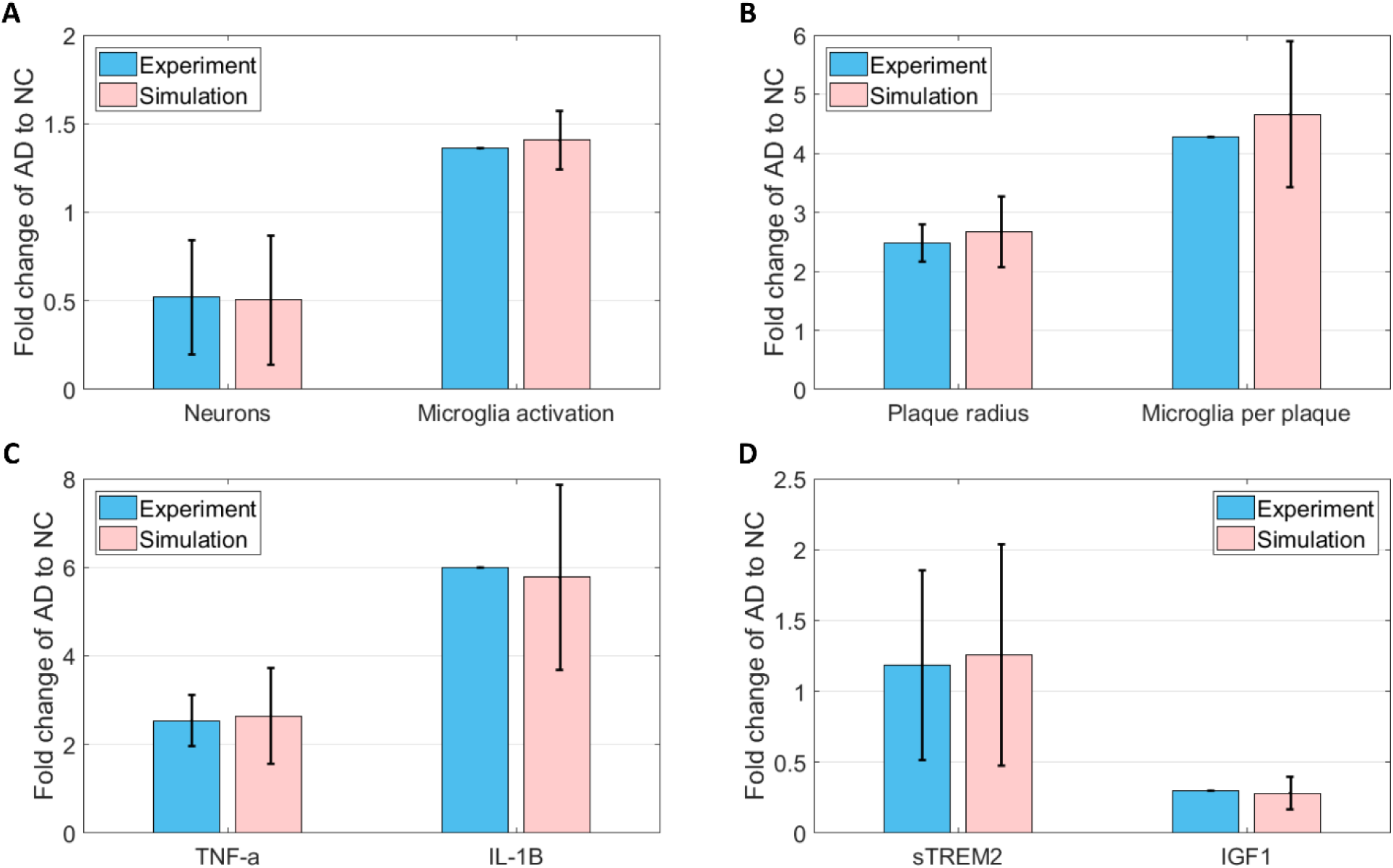
Model evaluation (data fitting). The experiment data were collected from literature (**Table 1**). In our simulation, 1 WK (Time step: 84) and 6 WK (Time step: 504) are considered as NC and AD condition, respectively.

To further validate the reliability of our MSMAD model, we compared the simulation results with the experimental data (**Table 2**) generated from 5xFAD mouse model [39]. Significantly more microglia are living in the AD mouse than wide type. We also confirmed increased expression of inflammatory cytokines (IL-1 β, and TNF-α) in 5XFAD mice. In summary, the experimental data further confirms that the outputs of MSMAD model are reliable.

**Table 2.** The experimental data used for model validation.

|  | WT | AD | Models | PMID |
| --- | --- | --- | --- | --- |
| Living microglia ($\times 10^5$) | $1.5 \pm 0.25$ | $3.8 \pm 0.45$ | 5xFAD mice | 25728668 |
| IL-1β (relative expression) | $1 \pm 0.3$ | $5.3 \pm 0.3$ | 5xFAD mice | 25728668 |
| TNF (relative expression) | $0.8 \pm 0.18$ | $2.16 \pm 0.47$ | 5xFAD mice | 25728668 |

### Prediction of potential therapeutic outcomes

To identify the potential therapeutic targets for AD progression in the brain microenvironment, we further simulated the effects of single or combine treatments (via anti-sTREM2, TREM2 agonist, and IGF-1 agonist) on neuron degradation and microglia phenotype switching using the established model. Anti-sTREM2 is assumed to inhibit sTREM2-induced pro-inflammatory signaling in microglia. TREM2 agonist is assumed to elevate TREM2 expression in microglia. IGF-1 agonist exerts neuroprotective effects. **Fig 5** shows the predicted neuron survival (**Fig 5A**) and microglia activation (**Fig 5B**) from various therapeutic strategies. For a single treatment, sTREM2 antagonist received the better effects than other two strategies in reducing neurodegeneration. In addition, TREM2 agonist has a better inhibitory effects on microglia activation than IGF1 agonist, which is consistent with the experimental observations shown in [31, 44]. Moreover, we found that the optimal prediction outcome was achieved from the treatment group with a combination of sTREM2 antagonist and IGF-1 agonist, indicating that inhibition of sTREM2 induced-pro-inflammatory signaling (e.g. NFKB pathway) and increase of IGF-1 concentration in AD brain potentially reduce neurodegeneration, as well as attenuate microglia activation. In the control condition (**Fig 5B**), the initial response of microglia was activated by accumulating Aβ plaques, and the late response might be triggered by increased sTREM2. Our simulations also show that the proportion of M1 and M2 microglia is significantly changed under different conditions (**Table S2**). Taken together, the combination of switching microglia phenotypes and the activation of insulin metabolism appear to impair AD development.

**Fig 5.**
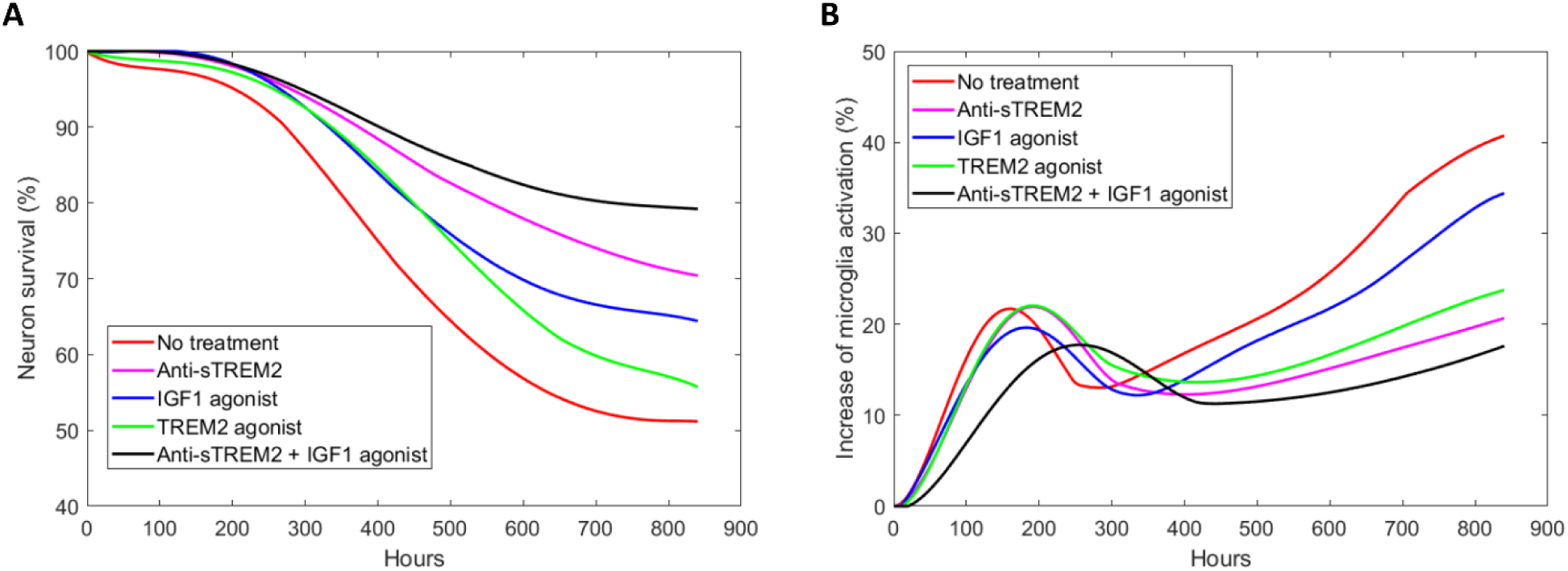
The effects of therapeutic strategies on neuron survival and microglia activation simulated by MSMAD model. The starting point of the curves is 1 WK (Time step: 84) in our *in silico* model. The curves presented are the mean effects of 100 runs of model.

## Discussion

To systematically understand the key role of microglia in AD progression and predict the optimal therapeutic strategy *in silico*, we developed a 3D multi-scale model (MSMAD) by integrating multi-level experimental data, to manipulate the neurodegeneration in a simulated system. We found that TRREM2/DAP12 signaling molecules, microglial phagocytosis related genes (CD68, Lgals3, etc.), and anti-inflammatory genes (e.g. IGF-1) in microglia were significantly up-regulated in the early stage of AD, but down-regulated in the late stage. In the meantime, the expression of NFKB pathway in microglia was elevated in the late onset of AD mouse, and further generated a series of pro-inflammatory factors, such as IL-1b, and TNF-*α*. Our analysis revealed that the microglia phenotypic switch appears to be driven by TREM2 signaling alteration. Switching of M1/M2 phenotypes of microglia may provide a new therapeutic perspective in AD treatment.

We are the first to systematically model the AD development using an integrated 3D system. In our MSMAD model, we simulated the neuron growth, microglia activation, and Aβ plaque formation (**Fig S2**). According to the timeline shown in [24], our simulation is up to 6 weeks, including the early stage (0-2 weeks) and late stage (2-6 weeks) of disease onset. The cell populations, the proportion of M1/M2 microglia, Aβ radius, and the density of microglia per plaque were calculated in each iteration (2 hours). Therefore, the proposed MSMAD model provides a new insight to simulate the dynamic changes in neuron degradation and immune response in AD mouse model. Current systems modeling framework also can be extended to human by fitting the model to AD patient-specific experimental data.

The MSMAD model also provides a novel computational platform to optimize the potential target therapy on AD progression. In our model, we assumed that anti-sTREM2 can block the downstream pathways of sTREM2, and inhibit the pro-inflammatory effects of microglia. The sTREM2 antagonist may inhibit the pro-inflammatory gene expression by modulating NFKB signaling. Since previous studies reported that sTREM2 levels is positively correlated with microglia activation and amyloidosis in PS2APP mice, reducing sTREM2 concentration in AD brain also seems to be a promising way to decrease AD risk [20]. TREM2 agonist is assumed to elevate the expression of TREM2 in microglia. Lee, *et al*. is the first to reveal that increasing TREM2 gene dosage reprograms microglia responsivity in AD mouse brains [31]. Different from the first two strategies, the aim of IGF-1 agonist is to directly elevate IGF-1 concentration, which might be beneficial to neuron survival. Recent studies indicates that intranasal insulin may ameliorate cognitive impairment, providing a potential way for treating AD patients [44, 45]. However, different subgroups of AD patients show different dose-response curves to intranasal insulin [46]. In the present study, we evaluated four new therapeutic strategies *in silico* with our optimized MSMAD model. The simulated results showed that optimal prediction outcome was achieved from the combination of sTREM2 antagonist plus IGF-1 agonist, revealing the important role of microglia phenotypic switch in AD development. Targeting any one of the vast pro-inflammatory mediators or pathways may not be efficacious in AD treatment.

Moreover, our MSMAD model includes a set of parameters, and most of parameters were tuned manually or determined based on the experimental results. To confirm the variability of the simulated results from the *in silico* model, parameter sensitivity analysis was performed by measuring the impact of small perturbation (5% increase) of individual 15 key parameters on neuron cell populations. We found that the 6^th^ and 8^th^ parameters (the basic rate of microglia proliferation, and the basic rate of M0 microglia change to M2) are more sensitive than others. The sensitivity analysis showed that the changes in model outcomes were under 4%, indicating that the established model were stable (**Fig S3**). In general, the model analysis provides us a strong belief on stability of the established model and model-based outcome prediction.

Finally, the quantified studies related with the proportion of M1 and M2 microglia activation were not reported. We are the first to evaluate the dynamic changes of microglial phenotypes in the AD progression using a novel computational model. Mastering the stage-specific switching of M1/M2 phenotypes at appropriate time may provide better therapeutic benefit.

## Methods

### Data collection and quantification

To optimize the parameters in agent-based model, we collected the experimental data related with the dynamic changes of cell population and cytokines from AD patients (or mouse models) relative to the normal controls (**Table 1**). Isla, et al. reported that the average total number of neurons in the cortex was reduced by 48% in the AD group [37]. Fan and colleagues evaluated the temporal profile of microglia activation in 30 subjects and found that microglia activation was increased by 36% in AD compared with controls [15]. Also, the mean plaque radius in the late-onset AD is significantly higher than that in early-onset [38]. Heslegrave, *et al*. examined CSF samples in 37 AD patients and 22 controls, and found that the sTREM2 concentrations were significantly higher in AD (p-value: 0.0457) [40]. Moreover, some pro-inflammatory cytokines (e.g. TNF-*α*, and IL-1β) were significantly increased in the late-onset of AD [41, 42]. In the meantime, recent experiments observed that IGF1 is significantly reduced in Alzheimer patients, which indicates that loss of IGF-1 input to the brain as an early biomarker of disease onset in AD [43]. For the microglia-plaque interactions, Wang, et al. observed that a high degree of microglial clustering around amyloid plaques in 5xFAD mice (average 4.28 microglia per plaque) [39].

### Multi-scale modeling of AD progression (MSMAD)

The multi-scale model developed in this study is to reflect the temporal imbalanced activation of M1/M2 microglia leading to AD progression. The proposed model combines a 3D multi-scale agent-based model (ABM) [47] for neurodegeneration and immune response, and a Hill functions system [48] for dynamic signaling transduction. The communications between them were mediated by cytokines, which were inferred from our genomic profiles. As a type of stochastic model, our ABM model used Markov Chain Monte Carlo approach [49] to simulate individual cell behaviors (**Fig S4**). Moreover, we also simulated the formation of Aβ plaques over time in our model based on the previous experimental observations [50]. The parameters in the ABM component should be tuned first to minimize the fitting error on the phenotype data shown in **Table 1**. Finally, based on the optimized model, we can design specific *in silico* therapeutic approaches to perturb the computational systems and predict the optimal intervention for preventing the development of AD. The details of MSMAD model as well as the corresponding mathematical rules was described in the **Supplementary Information**.

### Model implementation

The framework of ABM model was designed using the conception of “Object-Oriented Programming” and were achieved with C++. The proposed model was debugged and implemented under Linux environment on the cluster platform of the Texas Advanced Computing Center (TACC) at the University of Texas at Austin (http://www.tacc.utexas.edu). All of the parameters in the ABM model were tuned after running the system 100 times to fit the training data.

## Author contributions

XZ, ZJ designed the project. ZJ, CL analyzed the data. ZJ performed mathematical modeling and bioinformatics analysis. ZJ, CL, CS discussed results. ZJ and WZ wrote the manuscript.

## Conflicts interest

No potential conflicts of interest were disclosed.

## Funding

This work was supported by National Institutes of Health U01CA166886, U01AR069395-01A1 and R01GM123037 (Zhou). Funding for open access charge: Dr Carl V. Vartian Chair Professorship Funds to Dr Zhou from the University of Texas Health Science Center at Houston.

