## Supplementary information for "Systematically understanding the key roles of microglia in AD development"

### Supplementary Text. Development of MSMAD

#### 1.1 Simulated microenvironment and agents

In this study, we defined three types of cell agents in the MSMAD model to represent neuron, microglial cells, and AB. We initialized the AD brain mE as a 200×200×60 3D matrix (1 grid = 1 μm). Five types of small molecules diffuse and degrade in the mE, including Aβ from Neurons, IL-4 from M0 microglia, IGF-1 from M2 microglia, TNF-α, and IL-1β from M1 microglia. Aβ will deposit to form Aβ oligomer first, and followed by the accumulation of Aβ plaque. The initial numbers of neuronal and microglial cells are 10, and 20 respectively. Moreover, we randomly selected 10 points for the centers of Aβ plaque. The initial radiuses of these plaques are from 1 μm to 10 μm (1-10 grid), which are representative size reported by previous studies [1].

#### 1.2 The computational framework of MSMAD

Our MSMAD model is a hybrid model, which integrates Hill function system into agent-based model (ABM). The basic framework of ABM used in MSMAD was developed in our previous work [2]. The Markov Chain Monte Carlo approach was used to simulate cell behaviors of each individual cell [3]. As shown in **Fig S1**, cell behaviors were simulated by probability-based rule implementation. A cell sensed the cytokines/growth factors or drug doses from its neighborhood, processed them with signaling pathways (via Hill functions), and outputted the changes of probabilities of cell behaviors including cell proliferation rate, apoptosis rate, migration rate, and cytokine secretion rate. Cell decision was then determined by rolling a dice and compared with the probability of a given cell behavior. Details of cell behaviors for each type of cell agent as well as the corresponding rule was introduced in the following sections.

#### 1.3 Cell proliferation

Cell decision-making process was defined by agent rules, and the stochastic feature of the decision of an individual cell was realized by dice rolling simulation. Here, we used the cell decision-making of entering cell cycle as an example to elaborate the algorithm. Cell proliferation rules were defined according to proliferation rates with a dice $C_{rand} \epsilon [0,1]$. A cell cycle in our simulated system is 26 hours. If the dice fell into the interval [0, $P_{prol}$], the cell entered cell cycle and started to proliferate; otherwise, stayed quiescent. Four cell cycle phases (G0/G1, S, G2, and M) were defined [2]. Cells kept migrating during the first three phases, while tried to find a location to proliferate after entering the M phase. The duration of stage G0/G1 (10 hours) was determined by the Monte Carlo simulation, the S phase lasted for 6 hours (3 time steps), and the G2 and M phase would not exceed 6 hours (3 time step).

The proliferation rate of a neuron cell at location $(i, j, k)$ was described as shown in Eq. (1):

$$Growth\_N_{ijk} = NCgro0 * \left(1 + \frac{(\frac{IGF1_{ijk}}{K_{v1}})^n}{1+(\frac{IGF1_{ijk}}{K_{v1}})^n}\right) * \left(\frac{1}{1+(\frac{TNFa_{ijk}}{K_{v2}})^n}\right) \quad (1)$$

where $NCgro0$ is the initial proliferation rate of neuron. The proliferation can be promoted by IGF-1 and suppressed by TNF-a, respectively.

For microglia, we defined three conditions for representing M0, M2, and M1 phenotypes. The proliferation rate of a resting microglia (M0 phenotype) at location $(i, j, k)$ was defined in Eq. (2):

$$Prol\_M0_{ijk} = M0pro0 \quad (2)$$

In Eq. (2), $M0pro0$ is the initial proliferation rate of microglial cells. Considering the fact that increased AB density in local region may induce the activation of M0 microglia and transfer to M2 (initial immune response) [X], the probability of a M0 at location changes to M2 was shown in Eq. (3):

$$Prob\_M0\_M2 = tpchgpro0 * \left(1 + Fg_{m0} * A\beta_{ijk}\right) \tag{3}$$

In Eq. (3), tpchgpro0 is the basic rate for microglia changes from M0 and M2. $A\beta_{ijk}$ represents the $A\beta$ density in the local region (radius: 25 $\mu$m) relative to the location $(i, j, k)$. Increased $A\beta$ plaque may bind to TREM2 and promote M2 proliferation and survival:

$$Prol\_M2_{ijk} = M0pro0 * \left(1 + \frac{(\frac{A\beta_{ijk}}{K_{v3}})^n}{1+(\frac{AB}{K_{v3}})^n} + Fga_{m2} * D_1\right) \tag{4}$$

Where D1 represents the drug for elevating TREM2 expression [4]. After mid-term of AD, the protective role of microglia is lost and TREM2 expression is down-regulated. After that, sTREM2 concentration in CSF was significantly increased, indicating that M2 phenotype may be transformed to the M1 phenotype. sTREM2 potentially induces the pro-inflammatory effects of microglia by mediating the NFKB pathways [5]. Eq. (5) defines the probability of M2 phenotype transform to the M1 phenotype:

$$Prob\_M2\_M1 = typechgpro1\left(1 + Fgc_{m2} * (1 - D_2) * [\frac{\left(\frac{sTrem2}{K_{v4}}\right)^n}{1+\left(\frac{sTrem2}{K_{v4}}\right)^n}]\right) \tag{5}$$

sTREM2 also promotes microglia survival and proliferation in a PI3K/AKT-dependent manner [X]. The proliferation rate of M1 microglia was defined in Eq. (6):

$$Prol\_M1_{ijk} = M0pro0 * \left(1 + (1 - D_2)\frac{(\frac{sTrem2}{K_{v5}})^n}{1+(\frac{sTrem2}{K_{v5}})^n}\right) \tag{6}$$

Where D2 denotes the drug for inhibiting sTREM2-induced signaling.

### 1.4 Cell apoptosis

At each time step, if the apoptosis rate of a cell agent was lesser than the threshold, it would start apoptosis. Each cell takes 10 time steps to finish apoptosis and was then absorbed. The cell apoptosis is not only regulated by cell-death-related signaling pathways, but also is affected by the cytokines in the AD microenvironment.

The apoptosis rate of neuron is defined in Eq. (7):

$$Apop\_N_{ijk} = NCapop0 * \left(\frac{1}{1+\left(\frac{IGF1_{ijk}}{K_{v1}}\right)^n}\right) * \left(1 + \frac{\left(\frac{TNFa_{ijk}}{K_{v2}}\right)^n}{1+\left(\frac{TNFa_{ijk}}{K_{v2}}\right)^n}\right) * \left(1 + \frac{\left(\frac{A\beta_{ijk}}{K_{v3}}\right)^n}{1+\left(\frac{A\beta_{ijk}}{K_{v3}}\right)^n}\right) \tag{7}$$

Where $NCapop0$ denotes the initial apoptosis rate of neuron. According to Eq. (7), we can clearly see that the TNF-$\alpha$ and $A\beta$ oligomer induce neuron death, while IGF-1 exerts neuron protective effect. The apoptosis rate of M0 microglia is shown in Eq. (8):

$$Apop\_M0_{ijk} = M0apop0 \tag{8}$$

M2 microglia is activated as an initial response by $A\beta$ accumulation, and will have a transient reduction surrounding the amyloid plaque after clearance (Eq. (8)):

$$Apop\_M2_{ijk} = M0apop0 * [1 - Fgb_{m2} * D_1 + \left(\frac{1}{1+\left(\frac{A\beta_{ijk}}{K_{v3}}\right)^n}\right)] \quad (9)$$

For M1 microglia, previous studies reported that sTREM2 promotes microglia survival via PI3K/AKT pathways (see the following Eq. (10)).

$$Apop\_M1_{ijk} = M0apop0 * \left(1 + \frac{1}{1+(1-D_2)*\left(\frac{sTrem2}{K_{v5}}\right)^n}\right) \quad (10)$$

### 1.5 Cell migration

A non-M-phase cell at position *P* will migrate if it can find free space in its neighborhood (**Fig S1**). With a probability generated by dice rolling, a mature neuron generates a daughter cell at one of its adjoining locations. Neurons do not immediately form neural circuits through the growth of axons and dendrites. Instead, newborn neurons must first migrate long distances to their final destinations, maturing and finally generating neural circuitry. All the unoccupied positions ($P_{ijk}$) within a radius $r_{max}$ from the original position $P_0$ were scored and ranked [3]:

$$r_{max} = \left\lfloor \frac{2(1+4D*\Delta t)}{10} \right\rfloor + 1 \quad (11)$$

Where $D$ was the basic migration speed index, $\Delta t$ is the time step (2 hours). In addition, $p(r_l)$ was defined as the visiting chance of position $P_{ijk}$ with distance $r_l$ ($r_l \leq r_{max}$):

$$p(r_l) = \frac{1}{4\pi D\Delta t} \exp\left(\frac{{r_l}^2}{4\pi D\Delta t}\right) \quad (12)$$

Now, we firstly introduce the rules for NC migration. All the candidate unoccupied locations were ranked as Eq. (13):

$$R_l = p(r_l) * C_1 * C_2 * V_l \quad (13)$$

In Eq. (13), variable $C_1$ denotes if the candidate location $P_{ijk}$ exists an immediate neighbor as A β or M1 microglia. $C_2$ represents if there is an immediate neighbor of $P_{ijk}$ as a M2 microglia. The values of $C_1$ and $C_2$ were defined by Eq. (14-15).

$$C_1 = \begin{cases} 1.0, & \text{at least a neighbor is } A\beta \text{ or M1 microglia} \\ 0.10, & \text{otherwise} \end{cases} \quad (14)$$

$$C_2 = \begin{cases} 1.0, & \text{at least a neighbor is M2 microglia} \\ 0.20, & \text{otherwise} \end{cases} \quad (15)$$

In addition, $V_l$ described prostate cells try to avoid loneliness as well as crowdedness (see Eq. (16)).

$$V_l = \begin{cases} 1/8, & P_{ijk} \text{ has } 5-6 \text{ neighbor cells} \\ 1/4, & P_{ijk} \text{ has } 3-4 \text{ neighbor cells} \\ 1, & P_{ijk} \text{ has } 1-2 \text{ neighbor cells} \\ 1/16, & P_{ijk} \text{ has 0 neighbor cells} \end{cases} \quad (16)$$

After calculated the scores ($R_l$) of all the candidates, and then ranked them and selected the final location to migrate by dice casting (see the Eq. (17-18) in [2]).

Microglia cells migrate during CNS development and after CNS damage or disease. The migration ability regulated by their activation states. Previous findings indicate that M1 and M2 activated microglia differ in migratory and invasive capacity. Lively's work analyzed how these activation states affect microglia migration *in vitro* [6]. Their experimental observations revealed that M2 microglia exert stronger performance of migration than M1. The spatial ranges of microglia migration of M0, M1, and M2 were estimated from the data reported in [6]. A microglia cell can migrate from the original position $P_0$ to one of all the unoccupied positions ($P_{ijk}$) within a radius $r_{max}$. Here, the values of $r_{max}$ for M0, M1, and M2 are assigned as 6:2:12 (unit: grid). ). In addition, $p(r_l)$ was also defined as the visiting chance of position $P_{ijk}$ with distance $r_l$ ($r_l \leq r_{max}$). Here, we directly use the above Eq. (12).

Now, we firstly introduce the mathematical rules for microglia migration. All the candidate unoccupied locations were ranked as Eq. (13):

$$R_l = p(r_l) * C_1 * C_2 * C_3 * V_l \tag{17}$$

In Eq. (13), variable $C_1$ denotes if the candidate location $P_{ijk}$ exists an immediate neighbor as A β or M1 microglia. $C_2$ represents if there is an immediate neighbor of $P_{ijk}$ as a M2 microglia. $C_2$ represents if there is an immediate neighbor of $P_{ijk}$ as a M2 microglia. The values of $C_1$, $C_2$, and $C_3$ were defined by Eq. (18-19).

$$C_1 = \begin{cases} 1.0, & \text{if phenotype is M0, And at least a neighbor is A}\beta \\ 0.25, & \text{otherwise} \end{cases} \tag{18}$$

$$C_2 = \begin{cases} 1.0, & \text{if phenotype is M1, And at least a neighbor is NC} \\ 0.25, & \text{otherwise} \end{cases} \tag{19}$$

$$C_3 = \begin{cases} 1.0, & \text{if phenotype is M2, And at least a neighbor is NC or A}\beta \\ 0.25, & \text{otherwise} \end{cases} \tag{20}$$

In addition, $V_l$ described prostate cells try to avoid loneliness as well as crowdedness (see Eq. (21)).

$$V_l = \begin{cases} 1, & P_{ijk} \text{ has } 5-6 \text{ neighbor cells} \\ 1/4, & P_{ijk} \text{ has } 3-4 \text{ neighbor cells} \\ 1/8, & P_{ijk} \text{ has } 1-2 \text{ neighbor cells} \\ 1/16, & P_{ijk} \text{ has } 0 \text{ neighbor cells} \end{cases} \tag{21}$$

### 1.6 Dynamic distribution of cytokines

The dynamic 3D diffusion of growth factors (IGF-1, TNF-α, IL-1β, IL-4, etc.) in the simulated tumor microenvironment was defined as Eq. (22).

$$S_{ijk}(t+1) = (1 - DEG) * \left[S_{ijk}(t) * (1-\omega) + \frac{\omega}{6}\sum_{l=1}^{6} S_{ijk}^{l}(t)\right] \tag{22}$$

where $S_{ijk}(t+1)$ is the concentration of the factor on the position $(i, j, k)$ at the time step t+1, and $\omega$ is the diffusion constant. The value $S_{ijk}^{l}(t)$ indicates the concentration of a location, which is one of six immediate neighbors of position $(i, j, k)$. The constant $DEG$ represents the degrading rate of growth factors.

#### 1.7 AB plaque formation over time

Based on the previous experimental observations [1], we randomly initialized 10 points, which are considered as the centers for 10 Aβ plaques. The initial radiuses of these 10 Aβ plaques are from 1 to 10 μm. Each Aβ secreted from neuron will find the closest plaque and then gradually migrate toward it. The center point of each plaque will be updated each week (84 time steps). According to the experimental data shown in [1], we represented the dynamical changes of plaque radiuses (1-10 μm) over time with linear equations. All the formulas were listed as following:

$$\begin{cases} y = 1.94x - 0.94, & x \in [1,6], \text{initial size: } 1\mu m \\ y = 1.1x + 0.9, & x \in [1,6], \text{initial size: } 2\mu m \\ y = 1.2x + 1.8, & x \in [1,6], \text{initial size: } 3\mu m \\ y = 0.8x + 3.2, & x \in [1,6], \text{initial size: } 4\mu m \\ y = 1.44x + 3.56, & x \in [1,6], \text{initial size: } 5\mu m \\ y = 1.2x + 4.8, & x \in [1,6], \text{initial size: } 6\mu m \\ y = 1.1x + 5.9, & x \in [1,6], \text{initial size: } 7\mu m \\ y = x + 7, & x \in [1,6], \text{initial size: } 8\mu m \\ y = x + 8, & x \in [1,6], \text{initial size: } 9\mu m \\ y = 1.1x + 8.4, & x \in [1,6], \text{initial size: } 10\mu m \end{cases} \tag{12}$$

In this study, the time line defined in [7] was used in our simulations. Therefore, we just extracted the starting and ending points (early and late stage of AD) shown in the Figure 1D in the literature [1] to infer the above formulas (23). Except the above plaques, our model also simulate the appearance of newly formed β-amyloid plaques over time [1]. New plaques were mainly small size with 87% having a radius of < 4 μm.

### Supplementary Figures

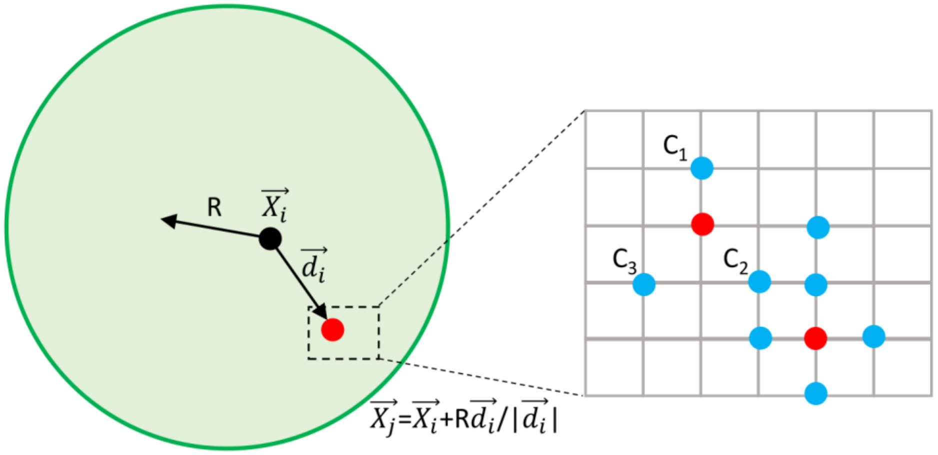


**Fig S1**. A sketch showing how spatial dispersal is implemented. A cell at position $X_i$ try to search all the candidate locations within the distance $R$, and found several empty positions (red dots). The probability $(M)$ of a cell moving from $X_i$ to $X_j$ was determined by: 1) the moving offset ($|d_i|$); 2) the number of occupied cells (blue dots) around the new position; and 3) which type of occupied cells is. In our model, $R$ equals 2 for migration, and 1 for proliferation.

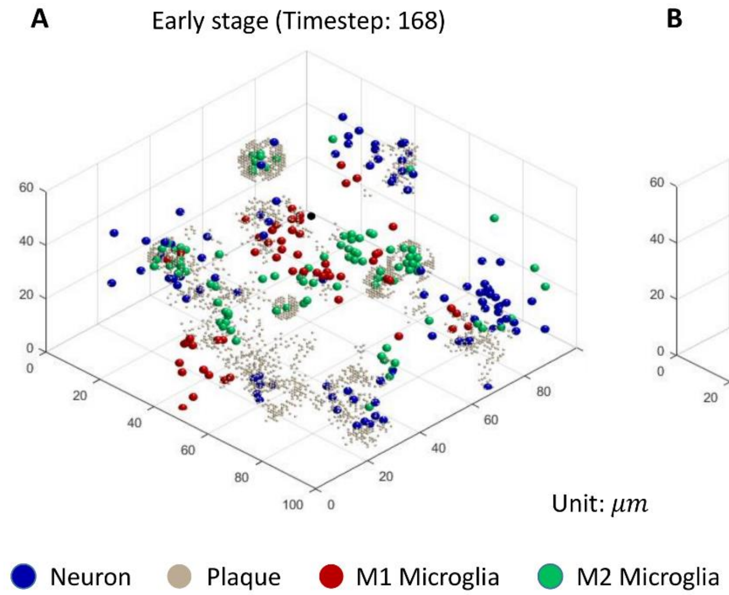


**Fig S2**. Aβ plaque formulation in the process of neurodegeneration. A representative case is presented.

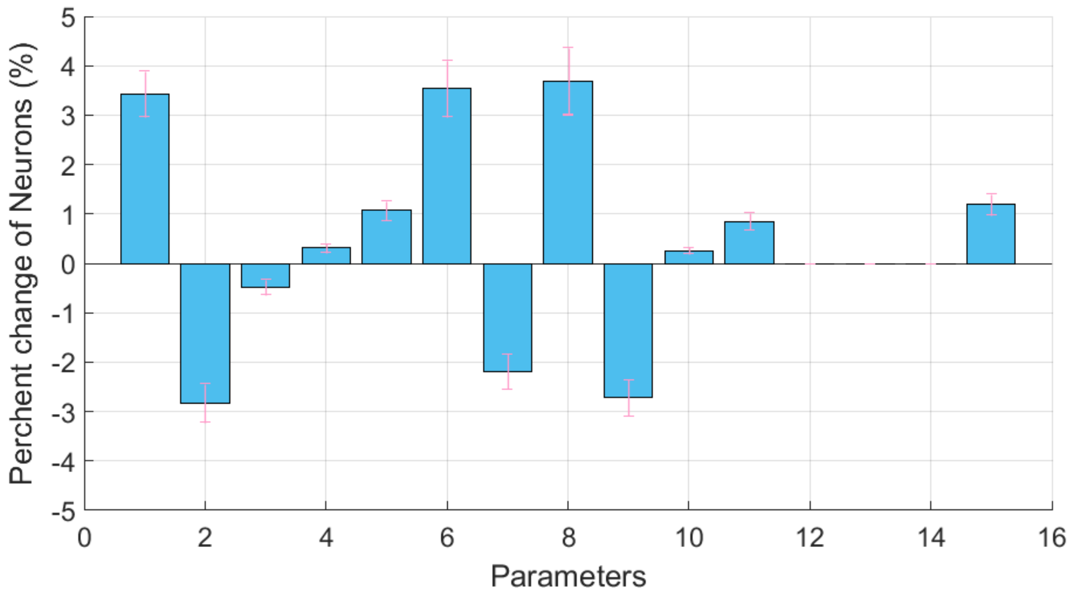


**Fig S3**. Sensitivity analysis of MSMAD model. The perturbation on each parameter is to increase 5%. Condition: no-treatment.

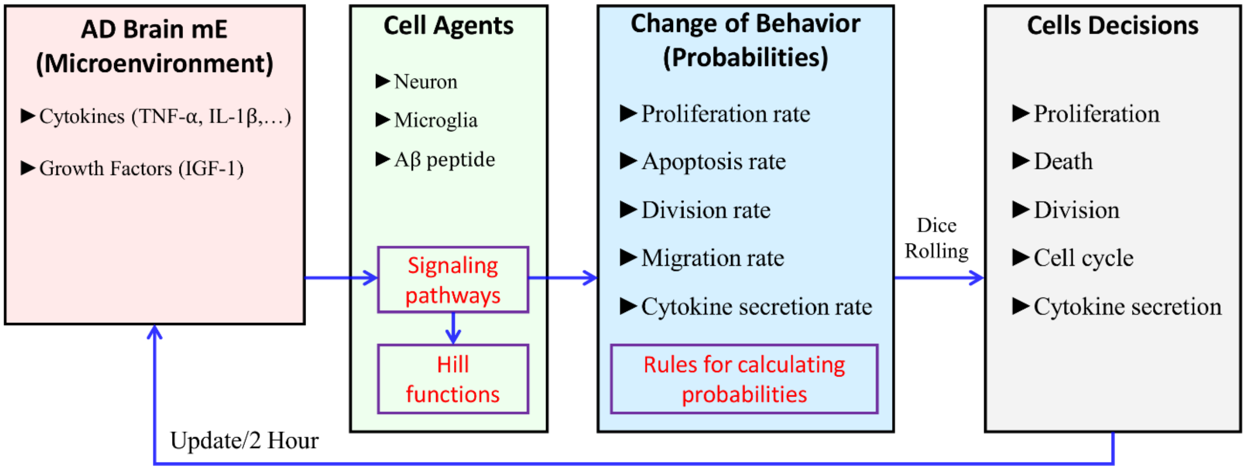


**Fig S4**. The stochastic simulation of cell behaviors.

### Supplementary Tables

**Table S1**. The estimated values of all the key parameters in ABM framework.

| No. | Variables | Descriptions | Values |
|---|---|---|---|
| 1 | $NCgro0$ | The initial proliferation rate of neurons | 0.225 |
| 2 | $NCapop0$ | The initial apoptosis rate of neurons | 0.021 |
| 3 | $K_{v1}$ | The coefficient of Hill Function in Eq. (1) described above | 0.0053 |
| 4 | $K_{v2}$ | The coefficient of Hill Function in Eq. (1) described above | 0.0050 |
| 5 | $K_{v3}$ | The coefficient of Hill Function in Eq. (7) described above | 0.08 |
| 6 | $M0pro0$ | The basic rate of microglia proliferation | 0.2753 |
| 7 | $M0apop0$ | The basic rate of microglia apoptosis | 0.0191 |
| 8 | $tpchgpro0$ | The basic rate of M0 microglia change to M2 | 0.056 |
| 9 | $tpchgpro1$ | The basic rate of M2 microglia change to M1 | 0.002 |
| 10 | $K_{v4}$ | The coefficient of Hill Function in Eq. (5) described above | 0.002 |
| 11 | $K_{v5}$ | The coefficient of Hill Function in Eq. (6) described above | 0.001 |
| 12 | $Fga_{m2}$ | The coefficient of the effect of TREM2 agonist on M2 growth | 0.3 |
| 13 | $Fgb_{m2}$ | The coefficient of the effect of TREM2 agonist on M2 death | 0.2 |
| 14 | $Fgc_{m2}$ | The coefficient of the effect of Anti-sTREM2 on inflammation | 1.5 |
| 15 | $Fg_{m0}$ | The coefficient of the effect of AB on transforming M0 to M2 | 1.0 |

**Table S2**. The prediction of M1/M2 proportion after single or combined treatment. $sTREM2^-$: anti-sTREM2. $IGF1^+$: IGF1 agonist. $TREM2^+$: TREM2 agonist (overexpression). A representative case was selected for each condition.

| | No treatment | $sTREM2^-$ | $IGF1^+$ | $TREM2^+$ | $sTREM2^-$ plus $IGF1^+$ |
|---|---|---|---|---|---|
| Early stage | 0.4524 | 0.224 | 0.411 | 0.365 | 0.176 |
| Late stage | 2.914 | 1.372 | 2.713 | 1.498 | 1.039 |
